# Not Just Noise: Aperiodic Brain Activity Reflects Corticospinal Excitability

**DOI:** 10.64898/2026.04.30.721880

**Authors:** Juliana R. Hougland, Miriam Kirchhoff, Timo van Hattem, Johanna Rösch, Jing Chen, Maxim Schaier, Paolo Belardinelli, Ulf Ziemann

## Abstract

**Background:** Electroencephalography (EEG) can be combined with transcranial magnetic stimulation (TMS) to perform brain-state-dependent stimulation. EEG-TMS studies have shown that corticospinal excitability, as measured via motor evoked potentials (MEPs), is modulated by pre-stimulus periodic EEG features, such as sensorimotor mu-rhythm phase and power. However, the influence of aperiodic brain activity on corticospinal excitability is largely unexplored.

**Objectives:** We evaluated the relationship between aperiodic and periodic mu-power, aperiodic exponent, and mu-phase on MEP amplitudes using EEG-TMS.

**Methods:** We applied 800 single TMS pulses to the left primary motor cortex in 78 healthy adults. We calculated aperiodic/periodic mu-power, aperiodic exponent, and mu-phase for each trial from the pre-stimulus C3-Hjorth transformed EEG. MEP amplitudes were extracted from the right first dorsal interosseous muscle. A linear mixed-effects model assessed relationships between MEP amplitudes and EEG features, with interactions between mu-phase and all other EEG features.

**Results:** Aperiodic and periodic mu-power, aperiodic exponent, and mu-phase significantly modulated MEP amplitudes. Higher aperiodic/periodic mu-power was associated with larger MEP amplitudes, while higher aperiodic exponent was associated with smaller MEP amplitudes. We found a significant interaction effect of aperiodic exponent and mu-phase on MEP amplitude. Aperiodic exponent was negatively associated with MEPs for trough, rising, falling phases, but positively associated with MEPs for peak phase.

**Conclusions:** Aperiodic and periodic features of brain activity are reflective of dissociable corticospinal excitability states. Future brain-state-dependent TMS interventions may include aperiodic EEG features, such as aperiodic mu-power and exponent, in addition to the well-established periodic features.

## Introduction

In recent years, substantial effort has focused on individualizing non-invasive brain stimulation. An increasingly popular method is brain-state-dependent stimulation, in which an individual’s endogenous brain activity is used to guide stimulation parameters (Bergmann, 2018; Schaworonkow et al., 2019; Wischnewski et al., 2022; Zrenner et al., 2018). Electroencephalography (EEG) is a convenient and temporally precise method for recording brain activity and can be combined with transcranial magnetic stimulation (TMS) to implement brain-state-dependent stimulation (Bergmann & Born, 2018; Zrenner et al., 2016; Zrenner et al., 2026). TMS induces an electric field in the brain via electromagnetic induction, which can lead to trans-synaptic activation of downstream neurons (Barker et al., 1985; Chen, 2000; Groppa et al., 2012; Siebner et al., 2022). When TMS is applied over the primary motor cortex (M1) at a suprathreshold intensity, the depolarization of descending corticospinal tract neurons can lead to a response in muscles of the contralateral limb, referred to as a motor evoked potential (MEP) (Barker et al., 1985; Rossini et al., 2015). MEP peak-to-peak volley amplitudes are widely used as an index of corticospinal excitability (Siebner et al., 2022; Spampinato et al., 2023).

Many brain-state-dependent stimulation protocols using EEG-TMS have primarily focused on utilizing features of the sensorimotor mu-rhythm (8-13 Hz). Multiple research groups found that the phase of the mu-rhythm oscillation is reflective of corticospinal excitability (Hussain et al., 2019; Ozdemir et al., 2022; Schaworonkow et al., 2018; Suresh & Hussain, 2023; Wischnewski et al., 2022; Zrenner et al., 2018; Zrenner et al., 2023). Larger MEPs are elicited when TMS is delivered at the trough/early rising mu-phase (high excitability state) versus the peak/early falling mu-phase (low excitability state). Moreover, long-term potentiation and long-term depression of corticospinal excitability have successfully been induced by delivering repetitive TMS to either the trough (potentiation) or peak (depression) of the ongoing mu-rhythm (Baur et al., 2022; Baur et al., 2020; Zrenner et al., 2018). Furthermore, researchers have investigated the association of other periodic EEG features with corticospinal excitability, such as mu-power, functional connectivity (e.g., mu-phase locking), and within other frequency bands (e.g., beta, gamma) (Haxel et al., 2025; Marzetti et al., 2024; Ogata et al., 2019; Ozdemir et al., 2022; Vetter et al., 2023; Wischnewski et al., 2022). However, current research has almost exclusively focused on periodic/oscillatory EEG features and overlooked the aperiodic component, or 1/f activity (**Figure 1**). In fact, studies evaluating the effects of mu-power on corticospinal excitability have either ignored the aperiodic component (Bergmann et al., 2019; Ozdemir et al., 2022; Thies et al., 2018) or removed it (Hussain et al., 2019; Wischnewski et al., 2022; Zrenner et al., 2023).

**Figure 1.**
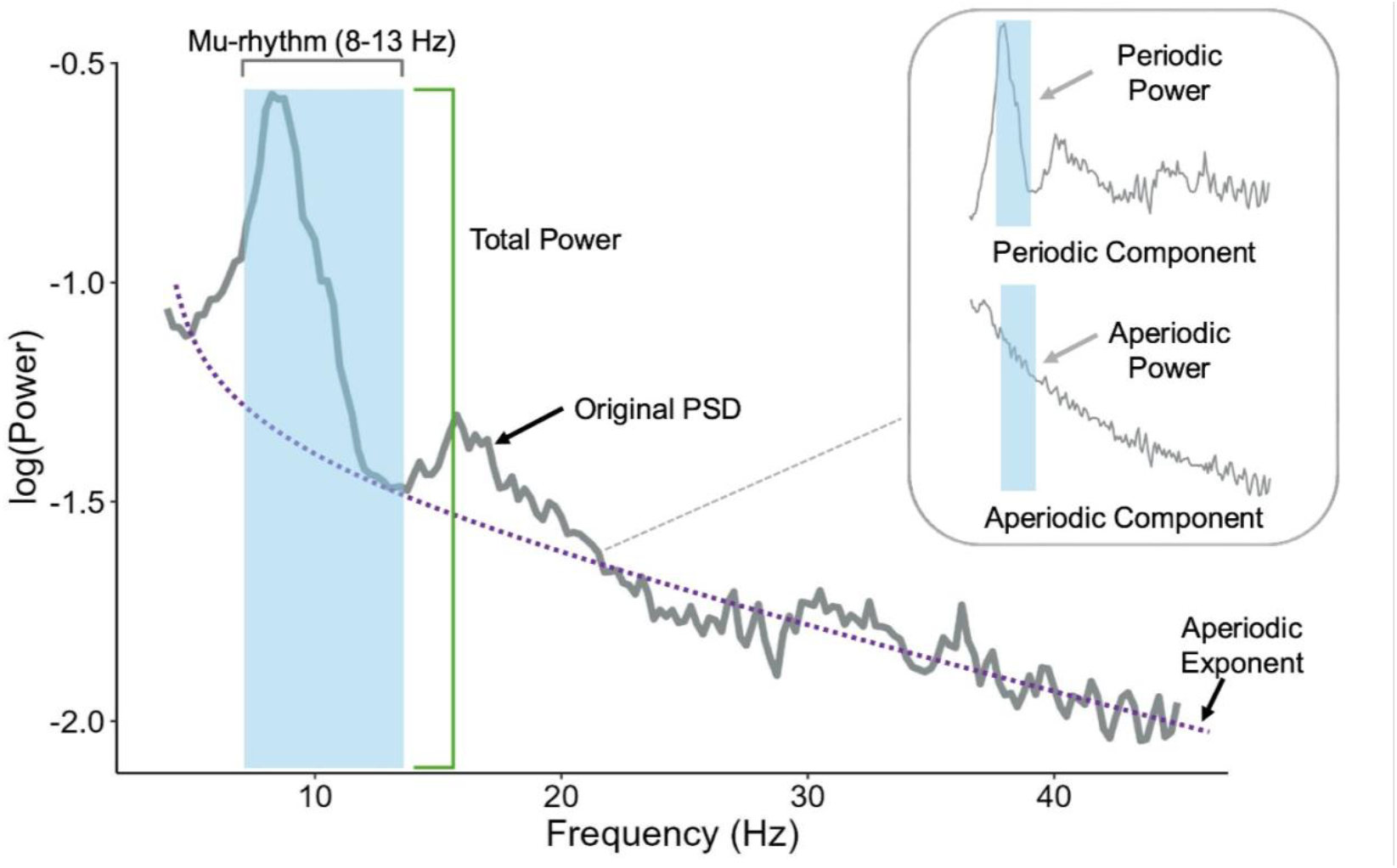
Periodic and Aperiodic Components of Neural Power Spectrum. The plot shows an example power spectrum. The grey line represents the full original power spectrum. The blue region highlights the mu-rhythm frequency band (8-13 Hz). The green bracket shows the total power within the mu-rhythm frequency band. The purple dotted line represents the aperiodic exponent or slope of the aperiodic activity. In the subpanel, the periodic (top) and aperiodic (bottom) components of the power spectrum are separated and the blue highlighted regions marks the mu-rhythm frequency band.

Neural power spectra can be separated into periodic and aperiodic components (**Figure 1**) through algorithmic modelling methods (Gerster et al., 2022). Periodic activity presents as narrowband peaks of power above the aperiodic component. These frequency-specific oscillations reflect behavioral and cognitive states (Buzsaki & Draguhn, 2004; Donoghue et al., 2020), and alterations in oscillatory features are implicated in various neurological and psychiatric disorders (Voytek & Knight, 2015). Although the aperiodic component was disregarded as background noise in the past, it has recently become a feature of interest. Aperiodic activity exhibits a 1/f-like distribution with decreasing power at increasing frequencies (Donoghue et al., 2020). The aperiodic component can be parameterized with a 1/f^*χ*^ function. The *χ*, or aperiodic exponent, represents the slope of the neural power spectrum, and has been implicated as a measure of physiological state. For example, the aperiodic exponent is believed to be a marker of the excitation/inhibition (E/I) balance (Gao et al., 2017; Voytek et al., 2015). A steeper slope indicates elevated inhibition, while a flatter slope suggests elevated excitation. The aperiodic exponent is altered in aging populations and those with neurodevelopmental and mental disorders such as autism, ADHD, and depression (Guirguis et al., 2025; Robertson et al., 2019; Sohal & Rubenstein, 2019; Voytek et al., 2015).

To date, only a few studies have explored the relationship between aperiodic brain activity and corticospinal excitability. Two studies have shown that the aperiodic exponent is inversely related to corticospinal excitability: as the exponent increases, MEP amplitudes decrease (Frohlich et al., 2024; Jin et al., 2025). Another study found that mu-rhythm aperiodic power is associated with a high excitability state (i.e., larger MEPs) (Jin et al., 2025). However, a third study found an interaction effect of aperiodic mu-power with mu-phase on MEP amplitudes, but not aperiodic mu-power alone (Suresh & Hussain, 2023).

This preliminary evidence therefore suggests that aperiodic brain activity reflects cortical excitability states and is a potentially valuable target feature for brain-state-dependent stimulation. Nevertheless, robust validation and replication of these initial results is needed. In the aforementioned studies, small subject and trial sample sizes may have limited the generalizability of the results (Frohlich et al., 2024; Jin et al., 2025; Suresh & Hussain, 2023). In the current study, we aim to analyze the relationship between aperiodic brain activity, including mu-rhythm aperiodic power and aperiodic exponent, and corticospinal excitability in a large dataset of 78 participants. Additionally, we looked at the largely unexplored interaction effects of aperiodic mu-rhythm power and aperiodic exponent with mu-phase on corticospinal excitability. In addition, we validated our findings in an independent dataset of 26 participants.

## Methods

### Participants

78 healthy, right-handed adults (42 females, 36 males, age = 24.4 ± 4.0 y, range: 19-36 y) completed one experimental session of EEG-TMS. All participants gave written informed consent and were excluded from the experiment if they had a history of neurological disease or substance use, or any contraindications to TMS (Rossi et al., 2021). This data overlaps with a dataset previously used by our group (Hougland et al., 2025; Zrenner et al., 2023). The study was approved by the Ethics Committee of the Medical Faculty at the University of Tübingen (Protocol number: 716/2014BO2) and conducted in accordance with the Declaration of Helsinki (2013 revision).

### Resting-state EEG, TMS and EEG-TMS

Participants were seated in a relaxed position, arms supported and elbows flexed at approximately 90 degrees, head position secured with a vacuum pillow (B.u.W Schmidt GmbH, Germany), and asked to fixate on a cross presented on a screen in front of them throughout all experimental procedures. First, a 10-minute eyes-open resting-state EEG measurement was recorded, using a 64-channel TMS-compatible cap (EasyCap, Germany), a 24-bit biosignal amplifier (NeurOne, Bittium, Finland), and a sampling frequency of 5 kHz. Impedances at all electrodes were kept < 5 kΩ throughout the session.

TMS was delivered to left motor cortex during EEG recording using a X100 stimulator and a Cool-B35 coil (MagVenture, Germany). Throughout TMS, neuronavigation was used to monitor coil location and orientation (Localite, Germany). The coil was held tangential to the scalp with an orientation of the coil handle 45° away from the midline. The biphasic current waveform induced with the second phase a lateral-posterior to medial-anterior current in the sensorimotor cortex. The TMS hotspot was identified as the scalp location over left motor cortex where the largest and most consistent MEPs were elicited in the first dorsal interosseous (FDI) muscle of the right hand (Rossini et al., 2015). The resting motor threshold (RMT) was defined as the stimulation intensity needed to elicit a MEP of at least 50 µV in 50% of trials (N = 58) (Rossini et al., 2015) or with the MT assessment tool, MTAT 2.0, which uses a maximum likelihood parametric estimation by sequential testing algorithm (N = 20) (Koponen & Peterchev, 2022). For the main experiment, 800 single TMS pulses were applied to the hotspot at 110% of RMT with an interstimulus interval of 2.25 ± 0.05 s (N = 58) or 2.0-3.0 s (N = 20). The MEPs were recorded with surface electromyography (EMG) from the right FDI, using a belly-tendon electrode montage.

### Preprocessing

All data were preprocessed and analyzed in MATLAB (v2025a, MathWorks, Natick, MA, USA) using the EEGLAB plugin (v2025.1.0), Python (v3.11.9, Python Software Foundation, Wilmington, DE, USA) and R (v4.2.2, R Foundation for Statistical Computing, Vienna, Austria).

Resting-state EEG: To determine each participant’s individual mu-peak frequency, the resting-state EEG data was downsampled to 1 kHz and a C3-Hjorth filter was applied (adjacent electrodes: FC5, FC1, CP5, and CP1) (Hjorth, 1975; Zrenner et al., 2023). The power spectral density (PSD) of the C3-Hjorth transformed resting-state EEG was calculated using Welch’s method (frequency range: 4-50 Hz, window length: 4 s, 50% overlap) and fit from 4-40 Hz using the FOOOF (fitting-one-over-f) algorithm (peak width limits: (1,8), maximum number of peaks: 6, minimum peak height: 0.2) (Donoghue et al., 2020). The mu-peak frequency was defined as the highest power peak frequency in the mu-frequency band (8-13 Hz). If no peak was identified, the mu-peak frequency was defaulted to 10 Hz (N = 15).

Pre-stimulus EEG: EEG-TMS data were epoched to 1000 ms pre-stimulus (-1005 to -5 ms) for each trial. Next, data were demeaned, linearly detrended, and downsampled to 1 kHz. A C3-Hjorth filter was applied, and trials exceeding the threshold range of 150 µV were excluded (Hougland et al., 2025; Kirchhoff et al., 2024; Zrenner et al., 2023). For mu-phase estimation, the data were filtered with an 80^th^-order FIR bandpass filter (stopband frequencies: mu-peak frequency ± 5, passband frequencies: mu-peak frequency ± 1) and the mu-phase of each trial at the time of the TMS pulse was estimated using the PHASTIMATE toolbox (filter order = 128, edge samples removed = 40, autoregressive model order = 20) (Zrenner et al., 2020). Trials were then binned into one of four equally spaced bins of width π/2 from – π to π: trough, rising, peak, and falling.

The PSD of the C3-Hjorth transformed trial EEG was calculated (frequency range: 4-50 Hz, window length: 1 s, 50% overlap), and the trial PSDs were separated into aperiodic and periodic components from 4-40 Hz using FOOOF (peak width limits: (2,6), maximum number of peaks: 4, minimum peak height: 0.1). The mean power in the mu-frequency band for the aperiodic and periodic components, the aperiodic exponent, and the goodness-of-fit (r^2^) were extracted for each trial. The aperiodic power, periodic power, and exponent were then z-scored. Trials with a goodness-of-fit below 5% (r^2^ < 0.05) were excluded. Additionally, we extracted the aperiodic exponent for each trial from each individual EEG electrode’s surface Laplacian-transformed PSD using the same parameters as above.

We chose to investigate only the mu-frequency band since it is the most commonly examined frequency band with regard to corticospinal excitability. Additionally, the aperiodic offset was not evaluated as our initial data analysis showed a very strong correlation with the aperiodic exponent and, therefore, provided no additional information.

EMG: EMG data were epoched 200 ms around the TMS pulse (-100 to 100 ms), downsampled to 1 kHz, baseline corrected, and linearly detrended. Line noise was removed from pre-stimulus EMG with a 50 Hz 4^th^-order Butterworth notch filter. MEP amplitudes were calculated as the peak-to-peak amplitude (maximum - minimum) of the FDI EMG from 20 to 40 ms post-stimulation. MEP amplitudes were natural log-transformed to reduce skewness of the distribution, and trials were excluded if pre-innervation (EMG amplitudes exceeding 50 µV in the pre-stimulus period) was present or the trial was identified as an outlier based on the 1.5 interquartile range rule.

### Statistical Analysis

A linear mixed-effects model (lme4 package) was used to assess the relationship between EEG features and corticospinal excitability. The response variable was the natural log-transformed MEP amplitudes. Fixed effects included z-scored aperiodic mu-power, z-scored periodic mu-power, z-scored aperiodic exponent, and mu-phase bin (four bins: trough, rising, peak, falling). Since interaction effects between mu-phase and mu-power have been previously reported (Hougland et al., 2025; Ozdemir et al., 2022; Suresh & Hussain, 2023), we additionally assessed interaction effects between mu-phase bin and the z-scored mu-powers (aperiodic and periodic) and aperiodic exponent. Subject was included as a random intercept. As a secondary analysis, we correlated the aperiodic exponent of each EEG electrode with that of C3 to assess whether the exponent is similar across different regions with the region of interest (left sensorimotor cortex). For each subject, we calculated the mean aperiodic exponent across all trials for each electrode, then correlated the mean aperiodic exponent of each electrode with C3 across all participants. Significance was set at *p* < 0.05 and post-hoc comparisons were corrected using the Benjamini Hochberg false-discovery rate (FDR) correction (Benjamini & Hochberg, 1995).

## Results

Mean RMT was 69 ± 10% of maximum stimulator output. Approximately 29% of all trials were excluded per subject. 6.6% were excluded for EEG noise threshold, 2.9% due to EMG pre-innervation, 10.9% due to MEP outlier, and 8.6% due to poor FOOOF fitting. In total, 43,622 trials were kept for final analysis.

### Effects of Aperiodic EEG on MEP Amplitude

Higher aperiodic mu-power was associated with larger MEP amplitudes (χ^2^(1) = 19.25, *p* < 0.001; see **Figure 2** for marginal model). Furthermore, there was a 5.5% increase in MEP amplitude per 1 SD increase in aperiodic mu-power. Aperiodic exponent was negatively associated with MEP amplitude (χ^2^(1) = 4.69, *p* < 0.05), such that a higher exponent reflected smaller MEPs (see **Figure 3A** for marginal model). A 1 SD increase in aperiodic exponent corresponded to a ∼2–3% decrease in MEP amplitude. The slope of the aperiodic exponent and MEP amplitude relationship for each subject can be seen in **Supplementary Figure 1**.

**Figure 2.**
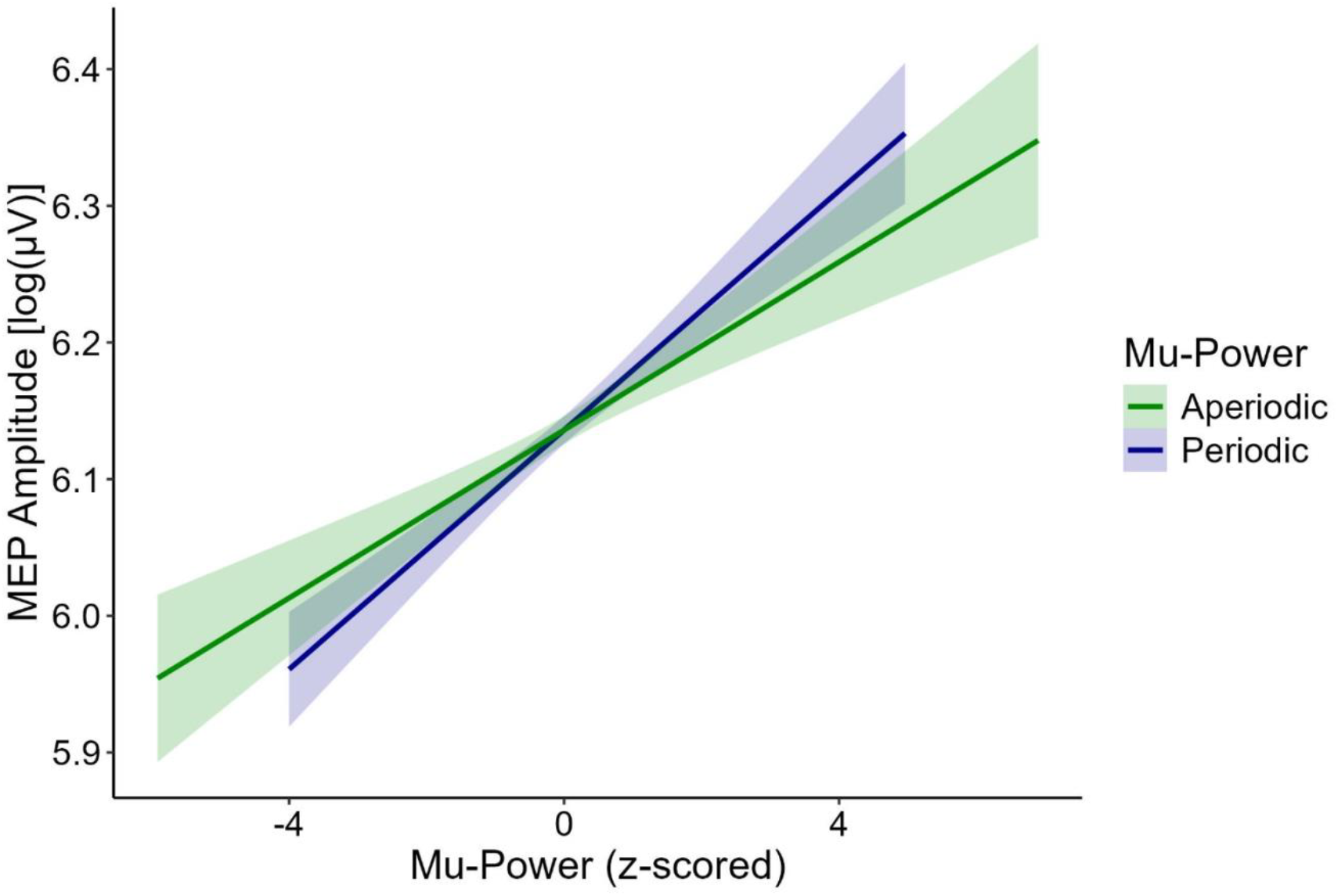
Effect of Aperiodic and Periodic Mu-Power on MEP Amplitude. The plot shows the influence of mu-power on natural log-transformed MEP amplitudes for both aperiodic (green line) and periodic (blue line) mu-power. Both lines represent the marginal linear model fit of mu-power and MEP amplitude and the shaded area is the 95% confidence interval. Aperiodic and periodic mu-power significantly modulated MEP amplitude. For both, higher mu-power reflected larger MEPs.

**Figure 3.**
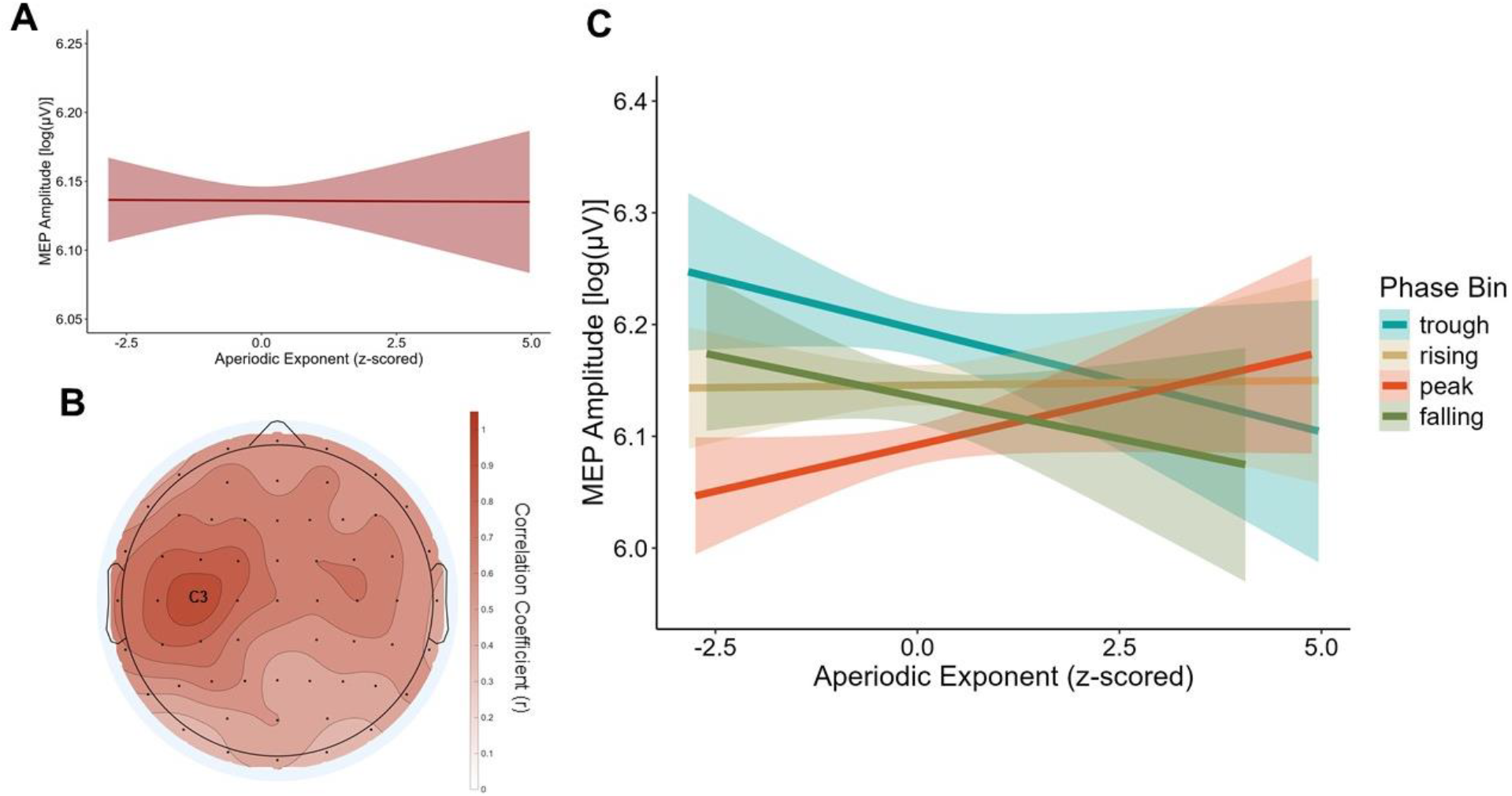
Effect of Aperiodic Exponent on MEP Amplitude. **A)** The plot demonstrates the marginal linear model fit of aperiodic exponent and natural log-transformed MEP amplitude. The shaded area reflects the 95% confidence interval. A significant effect of aperiodic exponent was found, in which a higher exponent was associated with smaller MEPs. **B)** The topography plot shows the correlation of the mean aperiodic exponent from each surface Laplacian-transformed EEG electrode with the surface Laplacian-transformed C3-electrode across all subjects and trials. The plot shows that aperiodic exponent is more correlated in sites near the C3-electrode but not in non-neighboring regions. **C)** The plot shows the influence of aperiodic exponent on natural log-transformed MEP amplitudes. Each line is color-coded by mu-phase bin and represents the marginal linear model fit of aperiodic exponent and MEP amplitude. The shaded are is the 95% confidence interval. A significant interaction effect was found between aperiodic exponent and mu-phase on MEP amplitude. The interaction effect on MEP amplitude was significantly different between peak and trough, and peak and falling phases.

### Spatial Specificity of Aperiodic Exponent Effects

To assess whether the aperiodic exponent reflects a global or local measure of excitability, we performed a correlation of the aperiodic exponent from each surface Laplacian-transformed EEG electrode with the aperiodic exponent of the surface Laplacian-transformed C3 electrode. The analysis showed local and some contralateral correlations of exponent with weaker correlations in non-neighboring electrodes (**Figure 3B**).

### Effects of Periodic EEG on MEP Amplitude

Periodic mu-power had a significant positive relationship with MEP amplitude, with higher power reflecting larger MEPs (χ^2^(1) = 22.49, *p* < 0.001; **Figure 2**). An increase of 1 SD in periodic mu-power was associated with 5.1% larger MEP amplitudes.

Mu-phase bin significantly modulated MEP amplitudes (χ^2^(3) = 78.06, *p* < 0.001; **Supplementary Figure 2**). The trough phase had the largest MEPs (6.19 ± 1.07 log(µV)), followed by rising (6.15 ± 1.09 log(µV)), falling (6.14 ± 1.10 log(µV)), and peak (6.09 ± 1.09 log(µV)). Moreover, trough MEPs were ∼9% larger than peak MEPs. Pairwise comparisons showed: 1. Trough MEPs were significantly larger than rising, peak, and falling MEPs (all FDR-corrected *p* < 0.001); 2. Rising MEPs were significantly larger than peak (FDR-corrected *p* < 0.001) but not falling (FDR-corrected *p* = 0.9225) MEPs; and 3. Peak MEPs were significantly smaller than all other phases (FDR-corrected *p* < 0.001).

### Interaction Effect of Mu-Phase with Mu-Power and Aperiodic Exponent on MEP Amplitude

MEP amplitudes were significantly modulated by the interaction of mu-phase and aperiodic exponent (χ^2^(3) = 13.66, *p* < 0.01), but not by the interactions of mu-phase with aperiodic (χ^2^(3) = 5.69, *p* = 0.128) or periodic mu-power (χ^2^(3) = 0.91, *p* = 0.823). Aperiodic exponent had phase-dependent effects on MEP amplitudes (see **Figure 3C** for marginal model). Trough, rising, and falling phase MEPs decreased with increasing aperiodic exponents, while peak phase MEPs increased. Simple slopes indicated that the negative relationship between aperiodic exponent and MEP amplitude was strongest during the falling phase (β = −0.040), moderate during trough (β = −0.027), minimal during rising (β = −0.008), and reversed during peak (β = 0.013). Pairwise comparisons indicated that the interaction effect of mu-phase and aperiodic exponent on MEP amplitude was significantly different between peak and trough (FDR-corrected *p* < 0.05), and peak and falling phases (FDR-corrected *p* < 0.01).

To assess the robustness of the interaction effect of mu-phase and aperiodic exponent on MEP amplitude, we performed additional linear mixed-effects models with 1. Z-scored aperiodic exponent added as a random slope; 2. Z-scored aperiodic exponent added as a random slope and mean periodic mu-power added as an interaction term; and 3. Z-scored aperiodic exponent added as a random slope and presence of a significant mu-phase effect on MEP amplitude (categorical: significant effect or insignificant effect) added as an interaction term. The mu-phase and aperiodic exponent interaction effect was present in all three models (all *p* < 0.05) indicating that the interaction effect is consistent across participants, and neither periodic mu-power nor significant mu-phase modulation of MEPs affects the mu-phase and aperiodic exponent interaction effect.

## Discussion

We analyzed the influence of aperiodic (aperiodic mu-power and exponent) and periodic (mu-phase and periodic mu-power) EEG features on corticospinal excitability measured with MEP amplitudes in healthy adults. Our results showed that aperiodic and periodic mu-power, aperiodic exponent, and mu-phase significantly modulated MEP amplitudes. Higher aperiodic and periodic mu-power were associated with larger MEPs, while higher aperiodic exponent was associated with smaller MEPs. Additionally, we presented novel findings on a mu-phase-dependent effect of aperiodic exponent on MEP amplitude. Overall, we found that aperiodic brain activity independently, and interactively with periodic brain activity, influences corticospinal excitability.

## Influence of Aperiodic Exponent on MEP Amplitude

The aperiodic exponent was, overall, negatively associated with MEP amplitude. The aperiodic exponent is representative of E/I balance; a higher exponent (steeper slope of 1/f activity) means relatively higher power at low frequency bands with a slow decay in power, which is characteristic of inhibitory GABA currents (Buzsaki et al., 2012; Donoghue et al., 2020; Gao et al., 2017). Thus, a high exponent is reflective of a high inhibitory state. As such, a higher aperiodic exponent should be indicative of smaller MEPs, which is the case in our and two other studies (Frohlich et al., 2024; Jin et al., 2025). In addition, we present novel findings investigating mu-phase-aperiodic exponent interaction effects on MEPs. The effect of aperiodic exponent was negative for trough and falling phases, slightly negative but almost flat for rising, and reversed (i.e., positive) at the peak phase, suggesting phase-dependent gating of E/I balance.

### Physiology of Phase-dependent Effects of Aperiodic Exponent on MEP Amplitude

Why would mu-phase alter the influence of aperiodic exponent on corticospinal excitability? Both the mu-rhythm oscillation and E/I balance are hypothesized to be modulated by two primary populations of neurons: excitatory pyramidal neurons and inhibitory interneurons (Klimesch et al., 2007; Lopes da Silva, 1991; Salinas & Sejnowski, 2000; Stein et al., 2026). Their interplay modulates both the mu-rhythm and E/I balance. However, if the same neuronal populations were contributing to both of these cortical activity features, we should expect to see the same influence of E/I balance (aperiodic exponent) on corticospinal excitability regardless of mu-phase. In contrast, our significant interaction results strongly suggest that mu-phase and aperiodic exponent reflect physiologically distinct excitability mechanisms.

The elevated corticospinal excitability at the trough of the mu-rhythm (Zrenner et al., 2018; Zrenner et al., 2020) is best explained by asymmetric pulsed facilitation (Bergmann et al., 2019). In contrast, pulsed inhibition (Jensen & Mazaheri, 2010; Klimesch et al., 2007) does not seem to play a role in determining these mu-rhythm phase-dependent corticospinal excitability states, as short-interval intracortical inhibition, a paired-pulse TMS-EMG metrics of GABAAergic inhibition (Ziemann et al., 1996) is not modulated by mu-rhythm phase (Bergmann et al., 2019). On the other hand, positive allosteric modulators at the GABAA receptor increase the aperiodic exponent (Salvatore et al., 2024), and MEP amplitude decreases under GABAAergic drugs (Di Lazzaro et al., 2000; Ziemann et al., 1996) and with increasing aperiodic exponent (present findings, and (Frohlich et al., 2024; Jin et al., 2025)). Moreover, decreasing spectroscopic concentrations of glutamate in human cortex are associated with an increasing aperiodic exponent (Sheldon et al., 2026), while antiglutamatergic drugs have no effect on MEP amplitude (Ziemann, Chen, et al., 1998; Ziemann, Tergau, et al., 1998), or even increase MEP amplitude (Di Lazzaro et al., 2003).

Our interaction data could then be interpreted in the following way: the trough of the mu-rhythm reflects a high-excitability corticospinal state characterized by high glutamatergic drive (pulsed facilitation). This high glutamatergic drive provides a high gain for activation of inhibitory interneurons (Molnár et al., 2008; Szegedi et al., 2016) so that increasing aperiodic exponent predominantly means increased GABAAergic inhibitory drive that in turn results in decreased MEP amplitude. In contrast, the peak of the mu-rhythm reflects a low-excitability corticospinal state characterized by low glutamatergic drive and low gain for activation of inhibitory interneurons. Increasing aperiodic exponent may then reflect predominantly decreasing glutamatergic activity, associated with increasing MEP amplitude. This interpretation is necessarily speculative and further experiments are required for substantiation, such as testing the effects of GABAAergic and antiglutamatergic drugs on the observed interaction of mu-phase with aperiodic exponent on MEP amplitude.

### Influence of Aperiodic and Periodic Mu-Power on MEP Amplitude

Both aperiodic and periodic mu-power had a significant positive relationship with MEP amplitude, indicating that higher mu-power is reflective of a higher corticospinal excitability state. Several studies that have examined mixed mu-power (combined aperiodic and periodic components) or periodic mu-power have demonstrated that higher power trials elicit larger MEPs (Karabanov et al., 2021; Ogata et al., 2019; Ozdemir et al., 2022; Suresh & Hussain, 2023; Thies et al., 2018). Nevertheless, these findings are conflicting, with some studies revealing a null or negative effect of mu-power on MEP amplitude (Karabanov et al., 2021; Madsen et al., 2019; Sauseng et al., 2009; Zarkowski et al., 2006). To our knowledge, only two studies have evaluated aperiodic power and corticospinal excitability (Jin et al., 2025; Suresh & Hussain, 2023). Similar to findings on periodic power, the studies showed conflicting results. In Jin et al. (2025), MEP trials sorted into the highest aperiodic alpha-power (C3-electrode only) group were significantly larger than MEPs in the low and lowest alpha-power groups (Jin et al., 2025). However, Suresh and Hussain (2023) found no influence of aperiodic mu-power on MEP amplitude, but did find a significant interaction effect with mu-phase, thus opposing the results of the current study (Suresh & Hussain, 2023). The reasons for these disparities are not fully clear. They may reflect biological variability in the tested populations and subtle difference in experimental setup and data analysis.

It is currently unclear if aperiodic and periodic power reflect different physiological processes. Our findings suggest that both aperiodic and periodic mu-power are very similarly indicative of corticospinal excitability, in which larger MEPs are elicited at higher power. Aperiodic and periodic power are additive components, which likely both contribute to the excitability state (Donoghue et al., 2020; Gerster et al., 2022). It has been suggested that the aperiodic component represents the baseline excitability state while periodic oscillations “fine-tune” excitability (Chini et al., 2022; Gao et al., 2017; Jin et al., 2025). Future work could explore potential physiological differences in aperiodic and periodic power by evaluating whether they are independently modulated by external stimuli such as action-observation, movement, or pharmacological intervention.

### Interaction Effects of Aperiodic and Periodic Mu-Power and Mu-Phase on MEP Amplitude

We did not find an interaction effect of aperiodic mu-power with mu-phase on MEP amplitude. Methodological differences in data acquisition and preprocessing could contribute to the difference in results with previous work. In the study from Suresh and Hussain (2023), which did see a significant interaction effect, mu-phase was targeted in real-time with only 25 trials per phase condition (trough, peak, and random) across 23 participants (Suresh & Hussain, 2023). Additionally, the study used a higher stimulation intensity of 120% of RMT, a longer interstimulus interval (5 s + random jitter), and a shorter pre-stimulus window (500 ms) for analysis (Suresh & Hussain, 2023).

Furthermore, there was no significant interaction effect of periodic mu-power and mu-phase on MEP amplitude, although previous studies have shown this effect (Hougland et al., 2025; Ozdemir et al., 2022; Suresh & Hussain, 2023). One reason for the null finding could be that our study’s sample has a large proportion of participants that do not demonstrate a strong mu-phase effect on MEPs. Although group-level mu-phase effects are common, multiple studies have reported that only approximately half of participants have a significant mu-phase effect (Hougland et al., 2025; Kirchhoff et al., 2024; Zrenner et al., 2023). In a previous study from our group, we found that if participants were split into two groups based on the presence of a mu-phase effect on MEPs, participants with a significant effect also showed a significant phase-power interaction, while participants with no effect only showed an effect of mu-power (Hougland et al., 2025). It is possible that the phase-power interaction is dampened in the current study due to a high proportion of participants without a significant mu-phase effect (∼59%).

### Influence of Mu-Phase on MEP Amplitude

Our findings on the effects of mu-phase on corticospinal excitability are well in-line with those of previous studies (Bergmann et al., 2019; Kirchhoff et al., 2024; Schaworonkow et al., 2019; Wischnewski et al., 2022; Zrenner et al., 2018; Zrenner et al., 2023). We showed a significant effect of mu-phase on MEP amplitudes, wherein TMS delivered at the trough phase elicited the largest MEPs while TMS at the peak phase elicited the smallest MEPs. Although a few studies have found no effect of only mu-phase on MEPs, the null effect is likely due to differences in data collection and analysis methods (Bigoni et al., 2024; Karabanov et al., 2021; Madsen et al., 2019; Ozdemir et al., 2022). Nevertheless, multiple independent research groups have shown that the trough and rising phases are representative of high excitability states (Suresh & Hussain, 2023; Wischnewski et al., 2022; Zrenner et al., 2018; Zrenner et al., 2023), which aligns with the results of the current study.

### Application for Brain-State-Dependent TMS

As mentioned previously, current brain-state-dependent TMS protocols primarily focus on mu-phase as an EEG feature of interest. However, mu-phase does not encapsulate total cortical excitability, and, by itself, does not explain high amounts of MEP variance (Kirchhoff et al., 2024). Multiple other EEG features are reflective of excitability, yet up to now, they have been largely ignored. In addition, phase is susceptible to real-time estimation errors due to narrowband filtering and data edge distortions (Jin et al., 2025; Liu et al., 2025; Zrenner et al., 2020). Aperiodic brain activity is one of these features that could have potential benefits for brain-state-dependent TMS application. The aperiodic exponent is a particularly useful feature as it alleviates the need for band-specific filtering or phase estimation (Frohlich et al., 2024; Jin et al., 2025). Moreover, algorithms that combine multiple features to deliver TMS at specific brain states may be advantageous for reducing variability in responses to TMS, and future research should be dedicated to the development and implementation of such protocols.

### Limitations

Our study has a few limitations. Firstly, there are multiple approaches to separating aperiodic and periodic components of brain activity. In the current study, we used the FOOOF algorithm since two of the other studies that assess aperiodic EEG and corticospinal excitability used this algorithm as well (Frohlich et al., 2024; Suresh & Hussain, 2023). However, other algorithms, such as IRASA (irregular-resampling autospectral analysis), could be used (Gerster et al., 2022; Wen & Liu, 2016). The two algorithms differ fundamentally in their approach: FOOOF uses Gaussian fitting to model the periodic components and a single aperiodic component, whereas IRASA just separates the periodic and aperiodic activity through a resampling procedure (Gerster et al., 2022). In practice, the two algorithms yield comparable results on clean, “easy” PSD data. However, differences in how both methods handle noisy PSD data could lead to divergent aperiodic component estimates (Gerster et al., 2022). Nevertheless, since future research would use aperiodic component fitting in real-time, brain-state-dependent settings, FOOOF has a practical advantage due to lower computational costs (Gerster et al., 2022). Additionally, short epochs of data limit the frequency resolution, which decreases the validity of aperiodic and periodic component separation. The intertrial interval varied between 2-3 s in the current study. To avoid influence of post-stimulus activity, we used a 1 s pre-stimulus interval for our analysis. Longer pre-stimulus epochs would help improve FOOOF fits for low-frequency oscillations.

## Conclusions

We showed that both aperiodic and periodic features of brain activity influence corticospinal excitability. In addition to the commonly investigated mu-phase effect on MEP amplitudes, MEPs increased with increasing aperiodic and periodic mu-power but generally decreased with increasing aperiodic exponent. Moreover, the effect of aperiodic exponent on corticospinal excitability was dependent on mu-phase. These findings suggest that aperiodic brain activity is not just background noise but reflects fluctuating cortical excitability states that can be dissociated from excitability states reflected by periodic EEG metrics. TMS interventions using brain-state-dependent stimulation should consider including aperiodic EEG features, such as mu-power and exponent, in addition to the well-established periodic features. Such protocols targeting both aperiodic and periodic EEG features could be beneficial for improving the efficacy of TMS interventions implemented for patient rehabilitation.

## Author Contributions

**Juliana R. Hougland:** Conceptualization, Investigation, Formal analysis, Visualization, Writing – original draft, Writing – review and editing. **Miriam Kirchhoff:** Formal analysis, Writing – review and editing. **Timo van Hattem:** Investigation, Writing – review and editing. **Johanna Rösch:** Writing – review and editing. **Jing Chen:** Investigation, Writing – review and editing. **Maxim Schaier:** Investigation, Writing – review and editing. **Paolo Belardinelli:** Writing – review and editing. **Ulf Ziemann:** Funding acquisition, Supervision, Writing – review and editing.

## Data Availability

Data will be made available upon request.

## Acknowledgements

This study is part of the ConnectToBrain project that has received funding from the European Research Council (ERC) under the European Union’s Horizon 2020 research and innovation programme (Grant agreement No. 810377).

