## Supplementary Materials for "Not Just Noise: Aperiodic Brain Activity Reflects Corticospinal Excitability"

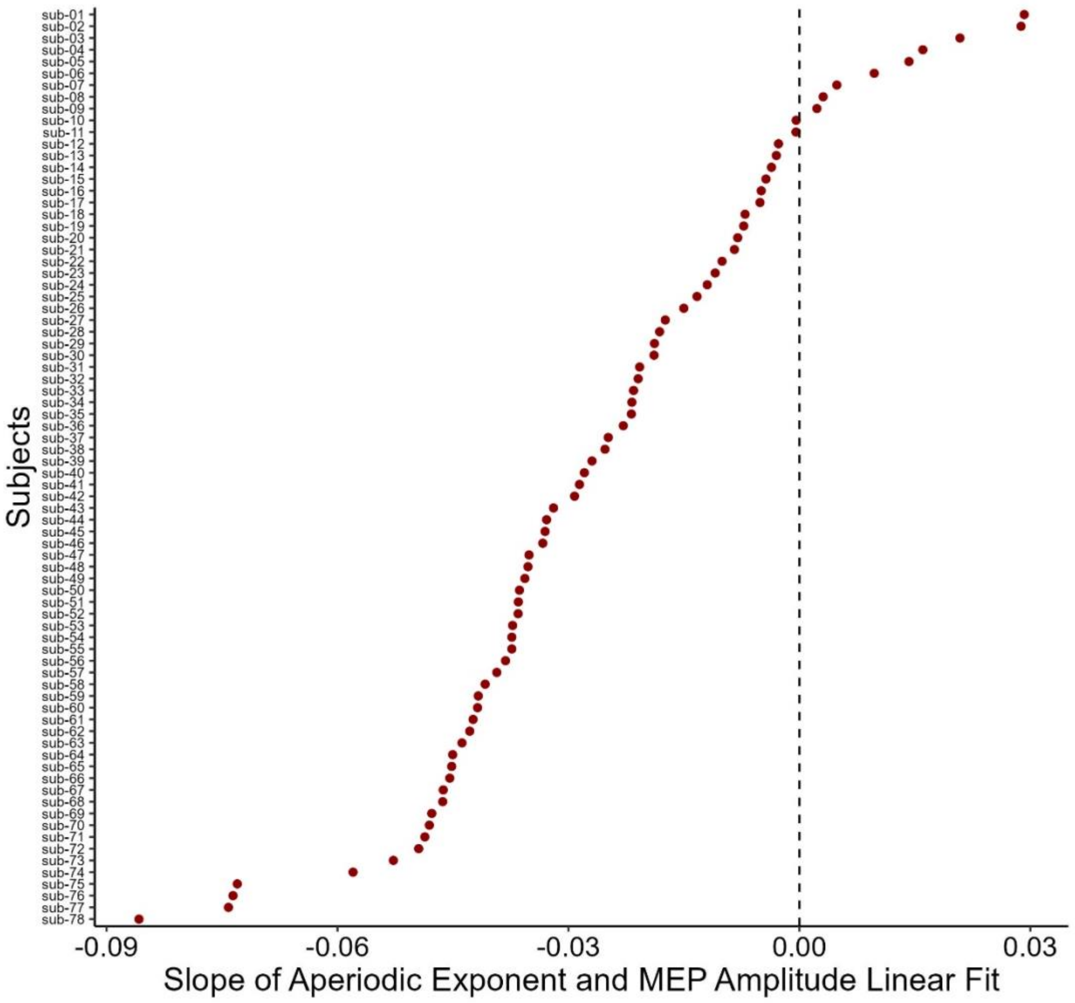


**Supplementary Figure 1. Slope of Aperiodic Exponent and MEP Amplitude Relationship Across All Subjects.** The plot shows the linear mixed-effects model random slope of aperiodic exponent for each individual participant. The dashed line is marked at zero to show where the relationship between aperiodic exponent and MEP amplitude changes from negative to positive. Only a few participants (~12%) had a positive relationship between aperiodic exponent and MEP amplitude, while the majority showed a negative relationship, indicating that a higher aperiodic exponent is associated with smaller MEP amplitudes.

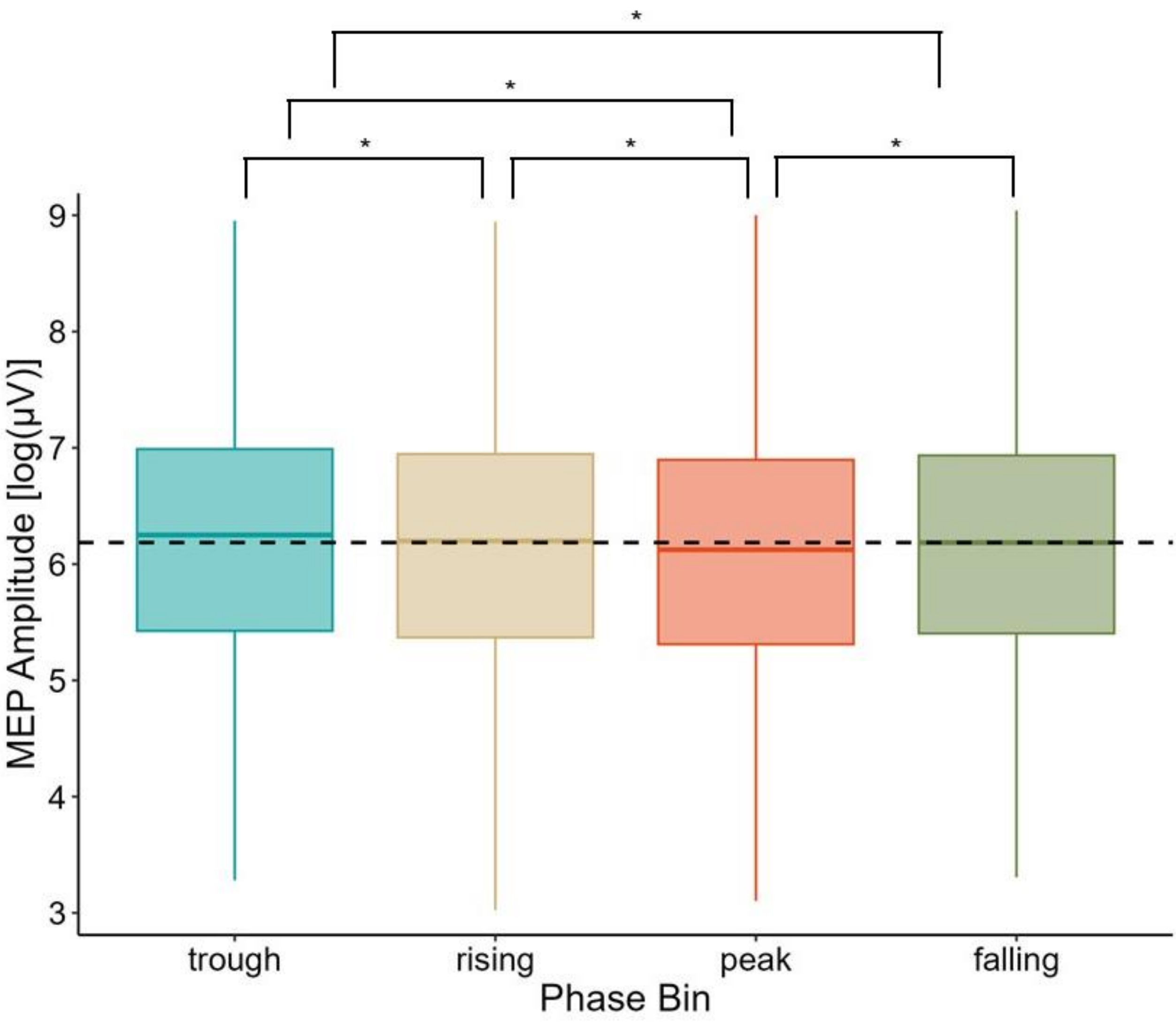


**Supplementary Figure 2. Effect of Mu-Phase on MEP Amplitude.** The boxplot displays the natural log-transformed MEP amplitudes across all subjects per mu-phase bin. The lower and upper hinges are first and third quantiles, and the lower and upper whiskers are 1.5 times the interquartile range of the corresponding hinge. The dashed line represents the median MEP amplitude across all phase bins. Results of the linear mixed-effects model showed a significant effect of mu-phase bin on MEP amplitude. Trough MEPs were the largest, followed by rising, falling, then peak MEPs. Trough MEPs were significantly larger than all other phases. Rising MEPs were significantly larger than peak but not falling MEPs, and peak MEPs were significantly smaller than all other phases. * FDR-corrected $p < 0.001$.
